# Exometabolomic analysis of decidualizing human endometrial stromal and perivascular cells

**DOI:** 10.1101/2020.08.06.221119

**Authors:** Sarah Harden, Jieliang Zhou, Maria Diniz-da-Costa, Emma S. Lucas, Liang Cui, Keisuke Murakami, Jinling Fang, Qingfeng Chen, Jan J Brosens, Yie Hou Lee

## Abstract

Differentiation of endometrial fibroblasts into specialized decidual cells controls embryo implantation and transforms the cycling endometrium into a semi-permanent, immune-protective matrix that accommodates the placenta throughout pregnancy. This process starts during the midluteal phase of the menstrual cycle with decidual transformation of perivascular cells (PVC) surrounding the terminal spiral arterioles and endometrial stromal cells (EnSC) underlying the luminal epithelium. Decidualization involves extensive cellular reprogramming and acquisition of a secretory phenotype, essential for coordinated placental trophoblast invasion. Secreted metabolites are an emerging class of signalling molecules. Here, we used liquid chromatography-mass spectrometry to characterise the dynamic changes in metabolite secretion (exometabolome) of primary PVC and EnSC decidualized over 8 days. We identified 79 annotated metabolites differentially secreted upon decidualization, including prostaglandin, sphingolipid, and hyaluronic acid metabolites. Secreted metabolites encompassed 21 metabolic pathways, most prominently glycerolipid and pyrimidine metabolism. Although temporal exometabolome changes were comparable between decidualizing PVC and EnSC, 32 metabolites were differentially secreted across the decidualization time-course. Further, targeted metabolomics demonstrated a conspicuous difference in xanthine secretion between decidualized PVC and EnSC. Taken together, our findings indicate that the metabolic footprints generated by different decidual subpopulations encode spatiotemporal information that may be important for optimal embryo implantation.

## INTRODUCTION

Cyclical decidualization, i.e. differentiation of endometrial fibroblasts into specialized decidual cells, is a hallmark of menstruating mammals. ^1,2^ Rather than being triggered by an implanting embryo, decidualization in menstruating species is initiated during the midluteal phase of each cycle in response to sustained progesterone signaling and rising intracellular cyclic adenosine monophosphate (cAMP) levels. ^3^ Initially, decidual changes are most prominent in sushi domain-containing 2-positive (SUSD2^+^) cells surrounding the terminal spiral arterioles and in SUSD2^-^ stromal cells underlying the luminal epithelium, ^4^ and then spread to encompass the entire stromal compartment. ^3^ In parallel, proliferating innate immune cells accumulate foremost uterine natural killer cells. ^5^ Upon embryo implantation, decidualizing cells rapidly encapsulate the conceptus, ^6,7^ engage in embryo biosensing and selection, ^8,9^ and then form an immune-privileged decidual matrix that controls interstitial and intravascular trophoblast invasion. ^3,10,11^

Recent studies have shown that decidualization is a multistep differentiation process, which starts with an evolutionarily conserved acute cellular stress response, ^12,13^ characterized by a burst of reactive oxygen species (ROS) production and release of proinflammatory cytokines. ^5,14–16^ After a lag period of several days, differentiating cells lose their fibroblastic appearance and emerge as secretory decidual cells with abundant cytoplasm and prominent endoplasmic reticulum. ^3^ At a molecular level, the decidual transformation of endometrial fibroblasts involves genome-wide remodeling of the chromatin landscape, ^17^ extensive reprogramming of multiple signal transduction and metabolic pathways, ^18–20^ and activation of decidual gene networks. ^21–23^ At a functional level, decidualization transforms endometrial fibroblasts into secretory cells that are anti-inflammatory, ^15,24^ resistant to stress signals, ^15,18,25^ and highly responsive to embryonic cues. ^8,26^

The decidual secretome is highly dynamic. Its composition changes across the different phases of the decidual pathway, ^15,16,27^ and the amplitude of the secretory response depends on the topography of cells in the native tissue. ^4^ For example, analysis of paired SUSD2^-^ and SUSD2^+^ cell cultures demonstrated that perivascular SUSD2^+^ cells secrete significantly higher levels of various chemokines and cytokines, including chemokine (C-C motif) ligand 7 (CCL7) and leukemia inhibitory factor (LIF). ^4^ These observations indicate that topographical microenvironments and chemokine gradients are established upon decidualization, which likely promote and direct trophoblast migration towards the spiral arterioles. Initially, invasion of the vessels results in plugging of these arteries by trophoblast. This enables the development of the conceptus in early pregnancy under low oxygen conditions, shielded from various environmental stressors. Notably, RNA-sequencing demonstrated that SUSD2^+^ stromal cells highly express genes encoding prototypic pericyte markers, including platelet-derived growth factor receptor β (*PDGFRB*), CD146 (*MCAM*), neural/glial antigen 2 (*CSPG4*), and α smooth muscle actin (*ACTA2*) ^28,29^. Hence, hereafter SUSD2^+^ cells are termed perivascular cells (PVC) whereas SUSD2^-^ stromal cells are referred to as endometrial stromal cells (EnSC). On average 6% and 94% of non-immune stromal cells isolated from midluteal biopsies are PVC and EnSC, respectively. ^27^

The metabolome encompasses a plethora of small organic molecules below 1.5 kDa with diverse physical and chemical structures, including nucleosides, amino acids, carbohydrates, and lipids. ^30^ Exometabolomics, also known as metabolic footprinting, is the study of how cells transform their surrounding microenvironment through the secretion of metabolites. Exometabolomics have been used to study the secreted metabolome profile of micro-organisms, with mammalian exometabolomics being a nascent field. ^31,32^ To date exometabolomics has been performed to understand differences in profile of breast cancer cell lines, T-cell immunity, and stem cells. ^33–35^

In this study, we used both untargeted and targeted LC-MS exometabolomics to map the dynamic changes in the metabolic footprints of 12 paired PVC and EnSC cultures decidualized over an 8-day time-course.

## MATERIALS AND METHODS

### Endometrial tissue collection

The study was approved by the NHS National Research Ethics – Hammersmith and Queen Charlotte’s & Chelsea Research Ethics Committee (1997/5065). Endometrial biopsies were obtained in the Implantation Clinic, a specialized research clinic at University Hospitals Coventry and Warwickshire National Health Service Trust. Written informed consent was given by all participants in accordance with The Declaration of Helsinki 2000. Endometrial biopsies, timed 6 to 10 days following the preovulatory luteinizing hormone surge, were taken using a Wallach EndocellTM sampler (Wallach, Trumbull, USA) following a transvaginal ultrasound scan to exclude overt uterine pathology. Further, none of the participants had been prescribed hormonal treatment in at least 3 months prior to biopsy.

### Isolation of PVC and EnSC from endometrial biopsies

Single-cell suspensions of EnSC were isolated from 12 midluteal biopsies as described previously. ^36^ In short, samples were collected in DMEM/F-12 with 10% dextran coated charcoal activated fetal bovine serum (DCC-FBS), finely chopped, and enzymatically digested with deoxyribonuclease type I (0.1 mg/mL; Roche, Burgess Hill, UK) and collagenase (0.5 mg/mL; Sigma-Aldrich, Gillingham, UK) for 1 h at 37°C. The cells were passed through a 40 µm cell strainer (Fisher Scientific, Loughborough, UK), which retained epithelial cells more resistant to enzymatic digestion. Ficoll-Paque PLUS (GE Healthcare, Little Chalfont, UK) was utilized to remove erythrocytes from the stromal cell fraction. Subsequently, SUSD2^+^ and SUSD2^-^ cells were isolated using magnetic bead sorting, as detailed previously. ^36^ Briefly, up to 1×10^6^ EnSC/100 µL of Magnetic Bead Buffer (0.5% BSA in PBS) were combined with 5 µL/1×10^6^ cells phycoerythrin (PE) conjugated anti-human SUSD2 antibody (BioLegend, London, UK). After 20 min on ice, approximately 1×10^7^ cells/80 µL of Magnetic Bead Buffer were incubated with anti-PE-magnetic-activated cell sorting MicroBeads (20 µL/1×10^7^ cells; Miltenyi Biotec, UK) for another 20 min. Cell suspensions (up to 1×10^8^ cells/500 µL of Magnetic Bead Buffer) were applied onto MS columns (Miltenyi Biotec) in a magnetic field, and washed multiple times with 500 µL of Magnetic Bead Buffer. SUSD2^-^ cells passed freely through the column, whereas magnetically labelled SUSD2^+^ cells were retained Following removal from the magnetic field, SUSD2^+^ cells were eluted from the columns with 1 mL of Magnetic Bead Buffer. Purified SUSD2^-^ and SUSD2^+^ cells were expanded in DMEM/F12 with 10% DCC-FBS, 1% antibiotic-antimycotic solution (Invitrogen), 1% L-glutamine (Invitrogen), estradiol (1 nM; Sigma-Aldrich), insulin (2 µg/mL; Sigma-Aldrich), and basic fibroblast growth factor (10 ng/mL; Merck Millipore, Watford, UK). Decidualization experiments were carried out at passage 2. Briefly, paired SUSD2^-^ and SUSD2^+^ cells were seeded 6-well plates at a density of 2×10^5^ cells/well and decidualized when 90% confluent with 0.5 mM 8-bromoadenosine cAMP (8-bromo-cAMP; Sigma-Aldrich) and 1 µM medroxyprogesterone acetate (MPA; Sigma-Aldrich) in phenol red-free DMEM/F12 containing 2% DCC-FBS, 1% antibiotic-antimycotic solution. The spent medium was collected every 48 h for analysis and the differentiation medium refreshed.

### Preparation of spent medium for analysis

For untargeted exometabolome analysis, samples were prepared as described previously, ^37,38^ with some modifications. Briefly 50 µL of EnSC and PVC conditioned media were thawed at 4°C, and quality control (QC) samples were prepared by mixing an equal amount of conditioned media from each of the samples. Both QC and the conditioned media from PVC and EnSC were processed in the same way. Proteins were precipitated with 200 µL ice-cold methanol containing 10 µg/mL 9-fluorenylmethoxycarbonyl-glycine as an internal standard (ISTD), and vortexed. Following centrifugation at 16,000 rpm for 10 min at min at 4 °C, the supernatant was collected and evaporated until dry in a speed vacuum evaporator then resuspended in 200 µL of water/methanol (98:2; v/v) for liquid chromatography-mass spectrometry (LC-MS) analysis. All samples were kept at 4°C and analyzed within 48 h. The sample run order was randomized to remove batch effects and the QC samples were analyzed after every eight samples to monitor the stability of the system.

### Untargeted LC-MS exometabolomics

Untargeted exometabolomics was performed as previously described.^39^ The prepared conditioned media was analyzed using Agilent 1290 ultrahigh-pressure liquid chromatography system (Waldbronn, Germany) equipped with a 6520 QTOF mass detector managed by a MassHunter workstation. The oven temperature was set at 45°C.

The 5 μL of injected sample was separated using an Agilent rapid resolution HT Zorbax SB-C18 column (2.1×50 mm, 1.8 mm; Agilent Technologies, Santa Clara, CA, USA), 0.4 mL/min flow rate, and a gradient elution involving a mobile phase consisting of (A) 0.1% formic acid in water and (B) 0.1% formic acid in methanol. To start the mobile phase was set at 5% B, with a 7-minute linear gradient to 70% B, then a 12 min gradient to 100% B. This was held for 3 minutes then returned to 5% B in 0.1 min.

Both positive and negative electrospray ionization were used to collect mass data between m/z 100 and 1000 at a rate of two scans per second. The ion spray was set at 4,000 V, and the heated capillary temperature was maintained at 350°C. The drying gas and nebulizer nitrogen gas flow rates were 12.0 L/min and 50 psi, respectively. Two reference masses (m/z 121.0509 (C5H4N4) and m/z 922.0098 (C18H18O6N3P3F24) were continuously infused to the system to allow constant mass correction during the run.

### Targeted LC-MS exometabolomics

Adenosine, inosine, hypoxanthine, and xanthine were measured and analyzed. The targeted LC-MS/MS analysis was conducted as described previously with some modifications ^34^. Briefly, LC-MS analysis was performed with Agilent 1290 ultrahigh-pressure liquid chromatography system (Waldbronn, Germany) coupled to an electrospray ionization with iFunnel Technology on an Agilent 6490 triple quadrupole mass spectrometer.

The auto-sampler was cooled at 4°C and 2 μL of injected sample was chromatographically separated using an Atlantis HILIC column (2.1×100 mm; Waters, Eschbornn, Germany). The mobile phases were (A) 10mM ammonium formate and 0.1% formic acid in water and (B) 0.1% formic acid in acetonitrile. Initially, 100% was utilized for 2 min, then reduced to 80% in a linear gradient for 11 min, and to 40% B over 1 min. This as held for 5 min and then the mobile phase was returned to starting conditions over 6 min. The column was kept at 45°C and the flow rate was 0.4 mL/min. Direct infusion of individual standard solutions allowed optimization of both the mass transition and collision energy for each compound by direct infusion. Both positive and negative electrospray ionization modes were performed with the following source parameters: drying gas temperature at 250°C with a flow of 14 L/min, sheath gas temperature at 400°C with a flow of 11 L/min, nebulizer gas pressure at 40 psi, capillary voltage 4,000 V and 3,500V for positive and negative mode respectively, and nozzle voltage 500 V for both positive and negative modes.

### Data analysis

Raw spectrometric data in untargeted metabolomics were analyzed by MassHunter Qualitative Analysis software (Agilent Technologies, US). The Molecular Feature Extractor algorithm was used to obtain the molecular features, employing the retention time (RT), chromatographic peak intensity and accurate mass as input. Next, the features were analyzed by MassHunter Mass Profiler Professional software (Agilent Technologies, US). Here, only features detected in at least 80% of the samples at the same sampling time point signal with an intensity ≥20,000 counts, three-times the limit of detection of the LC-MS instrument, were kept for further processing. The tolerance window for alignment of RT and m/z values was set at 0.15 min and 2 mDa respectively, and the data normalized to the 9-fluorenylmethoxycarbonyl-glycine ISTD spike.

Raw spectrometric data in targeted metabolomics were processed using MassHunter Workstation Quantitative Analysis software (Agilent Technologies, US). Missing values were obtained using half the lowest value. For both untargeted and targeted analysis, metabolites that were differentially expressed were inputted into the Metaboanalyst Pathway Analysis Tool to identify pathways that were altered during decidualization or between PVC and EnSC. This incorporates both pathway topology and enrichment analysis to output pathways that have changed significantly, and the pathway impact ^40^. The pathway impact is calculated by the number of compounds that are significantly altered in relation to the total number of compounds in the pathway. ^40^

### Statistical Analysis

For statistical analysis, nonparametric Test (Wilcoxon, Mann–Whitney test) with Holm-Sidak Multiple Testing Correction was employed and statistical significance was set at *p*<0.05. For multivariate data analysis using hierarchical clustering or partial least squares regression (PLSR) analysis, data were normalized by applying log2, median-centering and dividing by standard deviation. Hierarchical clustering unsupervised Euclidean distance hierarchical clustering was performed using HemI. ^41^ In addition, fold change (FC) analysis was performed to further filter the features and only those features with FC >1.5 were selected as potential significantly altered metabolites across decidualization.

### Compound identification

The identification of the differential metabolites structure was based on our published work. ^39^ Briefly, Masshunter software (Agilent) was used to calculate the elemental compositions of the metabolites based on their exact mass, the isotope pattern, and the nitrogen rule. The elemental composition in combination with the exact mass were used in open source database searches, including HMDB (http://www.hmdb.ca/), LIPIDMAPS (http://www.lipidmaps.org/), MassBank (http://www.massbank.jp/), and METLIN (http://metlin.scripps.edu/). Next, MS/MS experiments were performed to obtain structural information from the fragmentation pattern of the metabolite and these MS/MS spectra were searched and compared to compounds in the databases. Finally, the metabolites were confirmed by comparison with the standards where commercially available. The metabolites meet the minimum reporting standards for chemical analysis in metabolomics recommended by Metabolomics Standard Initiative (MSI). ^42^

## RESULTS

### Exometabolic footprints of decidualizing primary EnSC

The endometrial stromal fraction of 12 midluteal biopsies were separated into SUSD2^+^ PVC and SUSD2^-^ EnSC by magnetic-activated cell sorting (MACS). Both subpopulations were cultured to confluency and then decidualized with 8-bromo-cAMP and MPA (medroxyprogesterone acetate, a progestin) over 8 days. Conditioned culture media of paired decidualizing PVC and EnSC cultures were collected every 48 h and subjected LC-MS based metabolomics (Fig. 1). A total of 145 annotated secreted metabolites and 5 unidentified metabolites were detected in the conditioned media of both PVC and EnSC (Table S1). There were no metabolites unique to either PVC or EnSC. Hence, the metabolite data of PVC and EnSC were first combined to construct a temporal map of metabolic footprints associated with decidualization.

**Figure 1:**
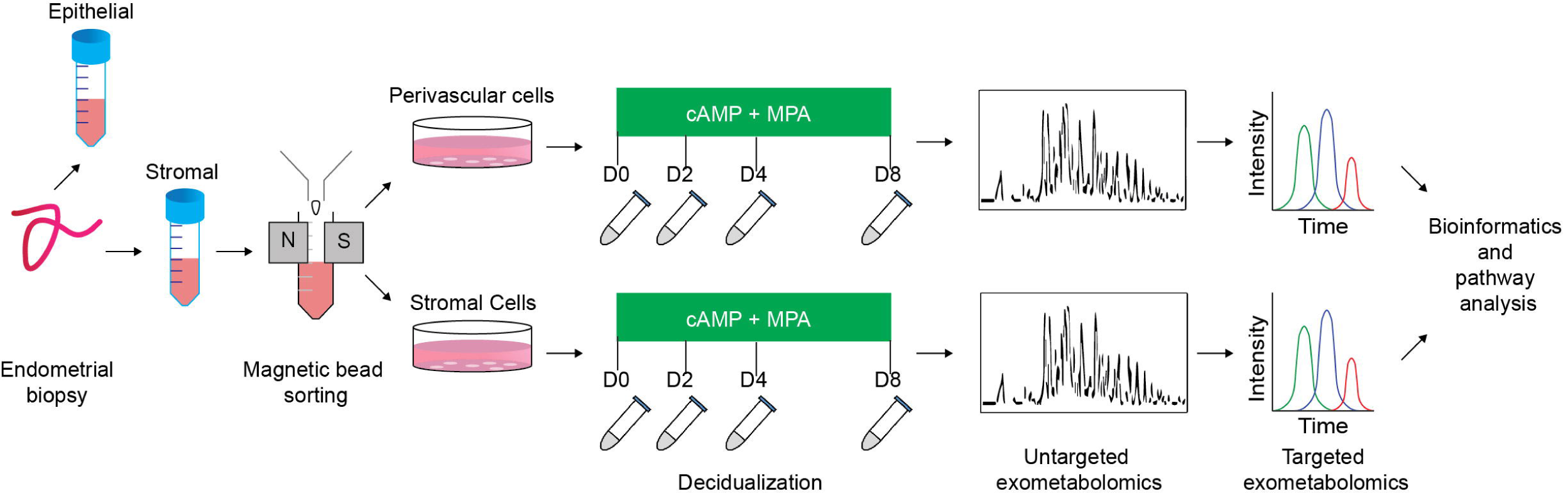
Schematic representation of experimental design and the analytical process. Paired PVC and EnSC cultured were established from 12 endometrial biopsies. Conditioned media was collected from undifferentiated cultures (D0, day 0) and after 2, 4, and 8 days of decidualization.

The QC samples clustered together in principal component analysis (PCA) scores plots (Fig. S1), indicating good stability and reproducibility of the chromatographic separation throughout the entire sequence. PLSR analysis separated the decidual time points into distinct groups (Fig. 2A). Regression coefficients were used to identify metabolites that shaped the progression of decidualizing cells throughout the time-course. Metabolites with the five highest β-coefficient (β) at each timepoint are depicted in Figure 2B. Phenylalanyl-tyrosine (β: 0.02), 2,3-dinor-8-isoprostaglandin F_2α_ (2, 3-dinor-8-isoPGF_2α_) (β: 0.01), and hypoxanthine (β: 0.02) were identified, respectively, as influencers on day 0 (undifferentiated cells), day 2 and day 4 of the decidual time-course. Adenosine thiamine triphosphate (AThTP) (β: 0.01) and hyaluronic acid (β: 0.01) were the highest influencers of day 8 of decidualization.

**Figure 2:**
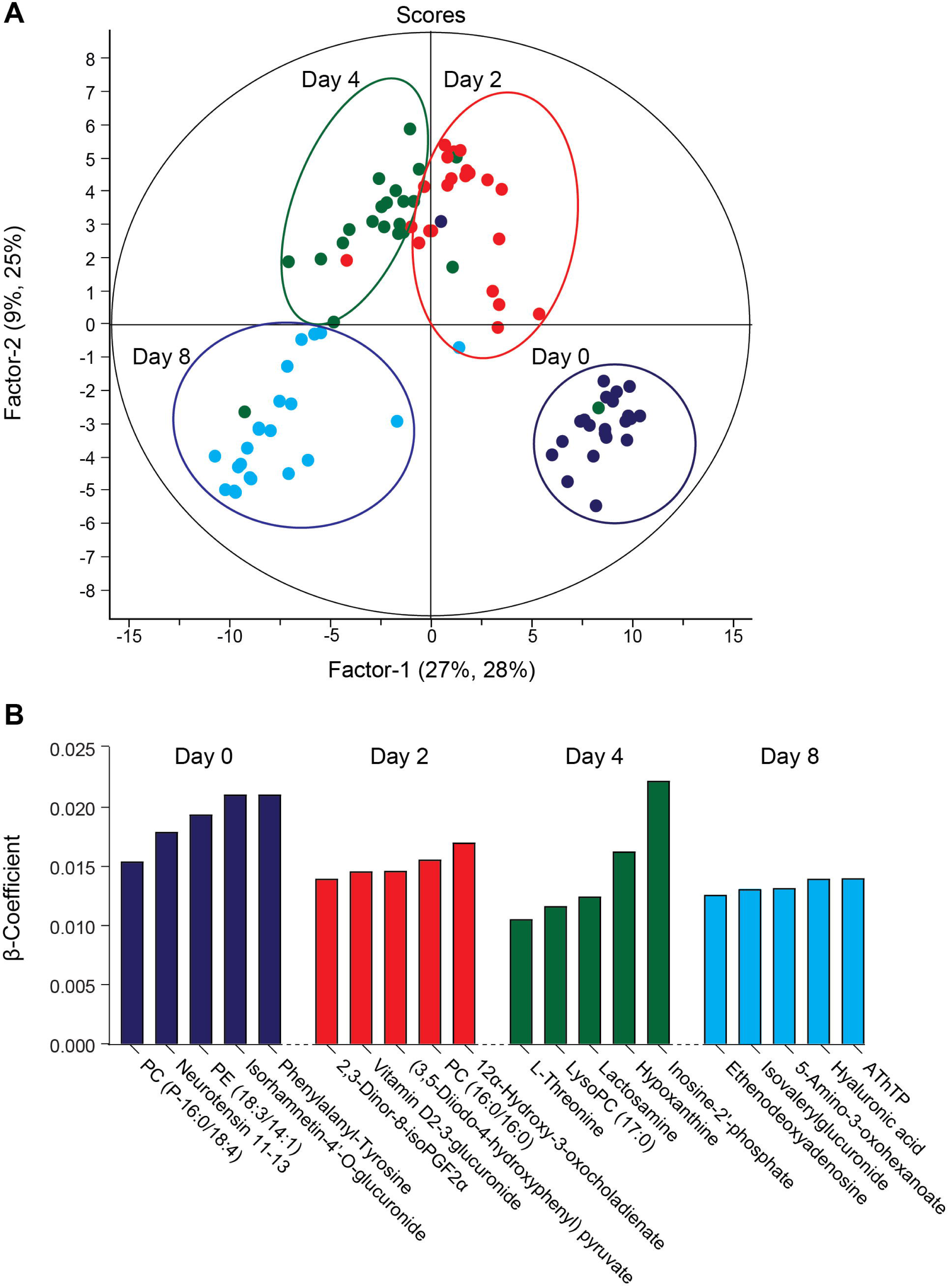
Temporal exometabolome of EnSC. A) Partial least squares regression (PLSR) analysis separated the exometabolomes according to decidual time-points. The secreted metabolite levels were log2 transformed, the data centred using median, and scaled by SD. B) Metabolites with the top five PLSR regression coefficients across the decidual time-course.

The exometabolome analysis of conditioned media revealed a marked temporal change in metabolic footprints upon decidualization. Out of the 150 detected compounds, the secreted levels of 79 metabolites changed by ≥1.5-fold (*p*<0.05) over the 8-day decidual time-course (Fig. 3A, Table S2). Unsupervised hierarchical clustering showed a major bifurcation in exometabolome profiles between undifferentiated and decidualizing cells. Further, each decidual time-point was characterized by a unique footprint, which supports the notion that decidualization is a multistep differentiation process (Fig. 3A).

**Figure 3:**
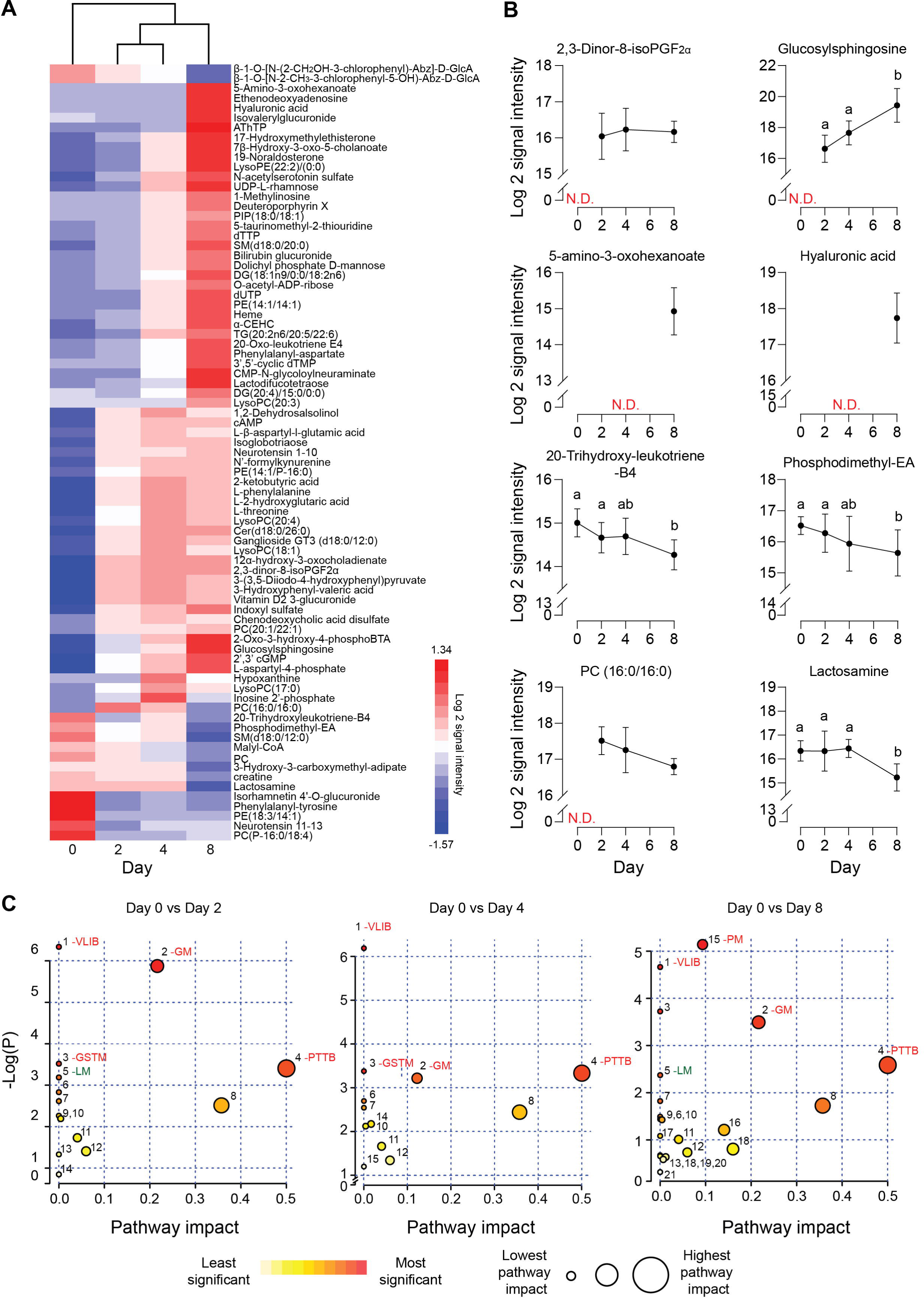
Exometabolic footprints of decidualizing primary EnSC. A) Heatmap of differentially secreted metabolites across the time-course based on unsupervised Euclidean distance hierarchical clustering. (B) Representative metabolites altered upon decidualization. Data represent mean ± SD. N.D, not detected. Different letters above the SD indicates differential secretion at the indicated time-point (*p*<0.05; *t*-test with Holm-Sidak correction). C) MetaboAnalyst pathway analysis across decidualization in EnSC. Enriched pathways include valine, leucine, and isoleucine biosynthesis (VLIB); glycero-phospholipid metabolism (GM); glycine, serine, and threonine metabolism (GSTM); Phenylalanine tyrosine and tryptophan biosynthesis (PTTB); Linoleic acid metabolism (LM); and pyrimidine metabolism (PM). Other pathways are tabulated in Supplementary Table 3.

Figure 3B highlights several regulated secreted metabolites, some of which are already implicated in decidualization. For example, the prostaglandin metabolite, 2, 3-dinor-8-isoPGF_2α_, which can be derived from PGE_2_, was undetectable in undifferentiated cultures but consistently upregulated in differentiating cells throughout the decidual time-course. Secreted levels of glucosylsphingosine, a sphingolipid metabolite, followed the same pattern, in keeping with the dependency of the decidual process on *de novo* sphingolipid synthesis. ^43,44^ Several metabolites were selectively secreted by the decidualizing cells on day 8, including 5-amino-3-oxohexanoate and hyaluronic acid (HA). HA prevents apoptosis of decidual cells through binding to its receptor CD44. ^45^ 20-trihydroxy-leukotriene-B4 and PE (18:3/14:1) are examples of metabolites inhibited upon decidualization, whereas phosphatidylcholine (PC) (16:0/16:0) and lactosamine exemplify metabolites exhibiting a biphasic secreted pattern, peaking on day 2 and day 4 of decidualization, respectively.

Metaboanalyst was used to identify metabolic pathways regulated upon decidualization. ^40^ Twenty-one metabolic pathways were identified across the decidual time-course (Table S3). As shown in Figure 3C, prominent pathways enriched across the decidual time-course included valine, leucine, and isoleucine biosynthesis (VLIB), glycerophospholipid metabolism (GM), and phenylalanine, tyrosine, and tryptophan metabolism (PTTM). Pyrimidine metabolism (PM) was enriched on day 8 of decidualization, indicating the requirement for pyrimidines, most likely as nitrogenous bases in DNA and RNA.

### Metabolites differentially secreted by decidualizing PVC and EnSC

The temporal changes in the metabolic footprints of decidualizing PVC and EnSC were comparable but not identical, as demonstrated PLSR analysis (Fig. S2). Notably, apart from a single metabolite (N2, N2-Dimethylguanosine), no significant differences were observed in the exometabolomes of undifferentiated PVC and EnSC. By day 2 of decidualization, however, the secreted levels of 12 annotated metabolites were significantly higher in PVC compared to EnSC and 4 were lower (≥1.5 fold-change, *p*<0.05; Table S4). By day 4 and day 8 of decidualization, secreted levels of 6 and 9 metabolites, respectively, were significantly different between PVC and EnSC (Table S4). Out of a total of 32 differentially secreted, annotated metabolites, 20 were more abundantly expressed by PVC. Figure 4A shows examples of metabolites differentially secreted between PVC and EnSC at each timepoint of the decidual pathway.

**Figure 4:**
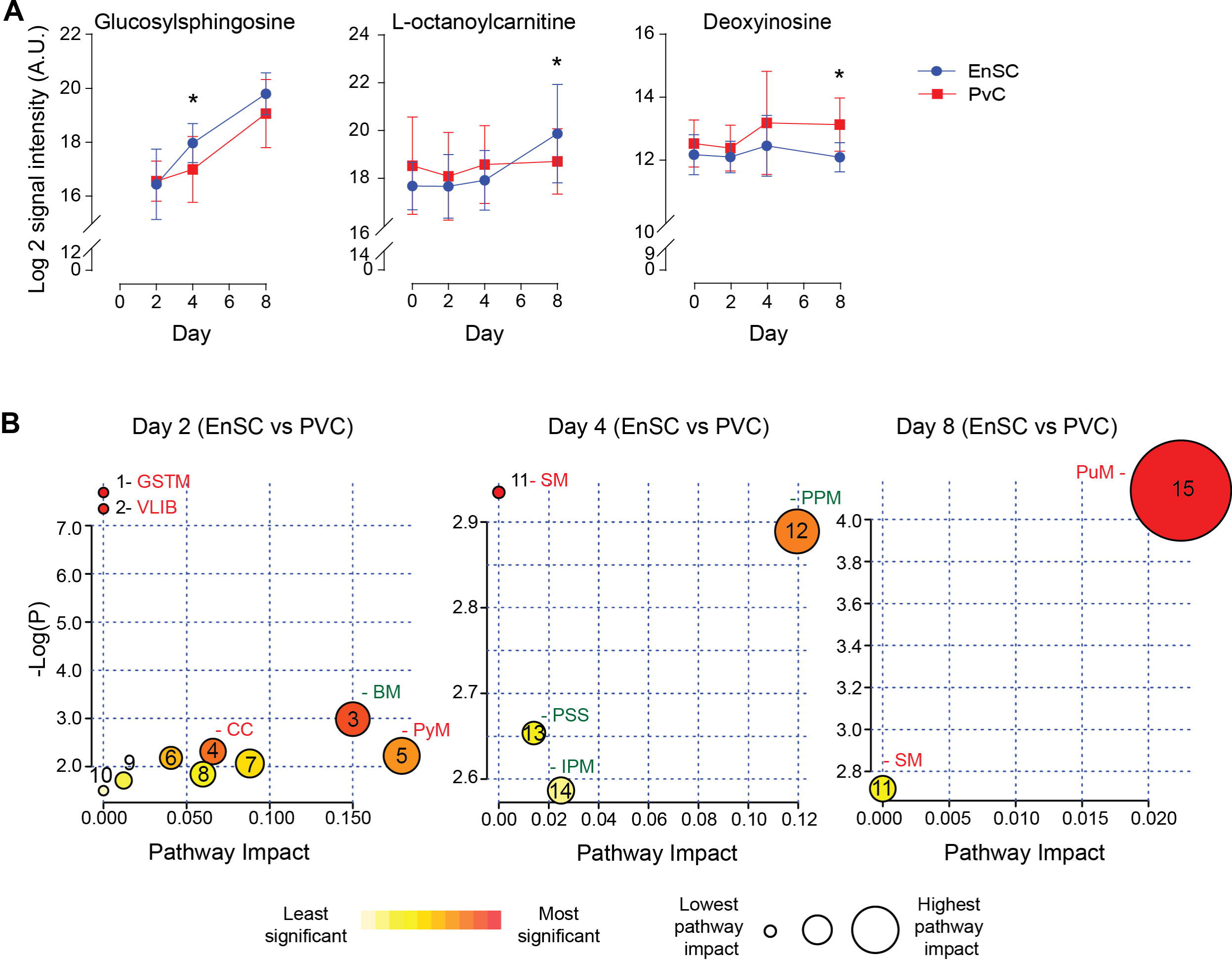
Metabolites differentially secreted by decidualizing PVC and EnSC. Data represents mean ± SD. * indicates *p*<0.05 (*t*-test with Holm-Sidak correction). B) Metaboanalyst Pathway analysis comparing PVC to EnSC of untargeted metabolomics by for day two, four and eight decidualization. Pathways labelled: glycine, serine, and threonine metabolism (GSTM), valine, leucine, and isoleucine biosynthesis (VLIB), biotin metabolism (BM), citrate cycle (CC), pyruvate metabolism (PyM), sphingolipid metabolism (SM), pentose phosphate metabolism (PPM), phosphatidylinositol signalling system (PSS), inositol phosphate metabolism (IPM) and purine metabolism (PuM). Other pathways can be found in Supplementary Table 5.

To further elucidate the metabolic differences between PVC and EnSC upon decidualization, pathway enrichment analysis was performed (Fig. 4B). Interestingly, distinct metabolomic profiles emerged. The citric cycle (CC) cycle, and glycine, serine and threonine metabolism (GSTM) were enriched in PVC cells on day 2 of decidualization. Enrichment of sphingolipid metabolism (SM) arose at in PVC at day 4, whereas inositol phosphate metabolism (IPM) and the pentose phosphate metabolism (PPM) were more prominent in EnSC. Finally, on day 8, sphingolipid (SM) and purine metabolism (PuM) were enriched in PVC. Purine metabolism had the highest pathway impact and significance (Fig. 4B), suggesting a greater autocrine or paracrine requirement of purine signaling upon decidualization in PVC when compared to EnSC.

### Targeted metabolomic analysis

The enhancement in purine metabolism prompted us to investigate their wider metabolomic network in greater depth. We developed an analytically robust targeted mass spectrometry analysis (average coefficient of variation=23.9%) to assess adenosine, inosine, hypoxanthine, and xanthine levels. Adenosine (7.8 fold-change, *p*=0.0001), inosine (1.53 fold-change, *p*<0.02) and xanthine (3.1 fold-change, *p*=0.0001) were significantly higher in PVC on day 8 of the decidual time-course in comparison to undifferentiated cells whereas uridine (0.21 fold-change, *p*<0.0001) and cytidine (0.42 fold-change, *p*=0.001) were significantly lower. Furthermore, xanthine levels were significantly lower in PVC compared to EnSC conditioned media in both undifferentiated and decidualized cultures (Fig. 5B).

**Figure 5:**
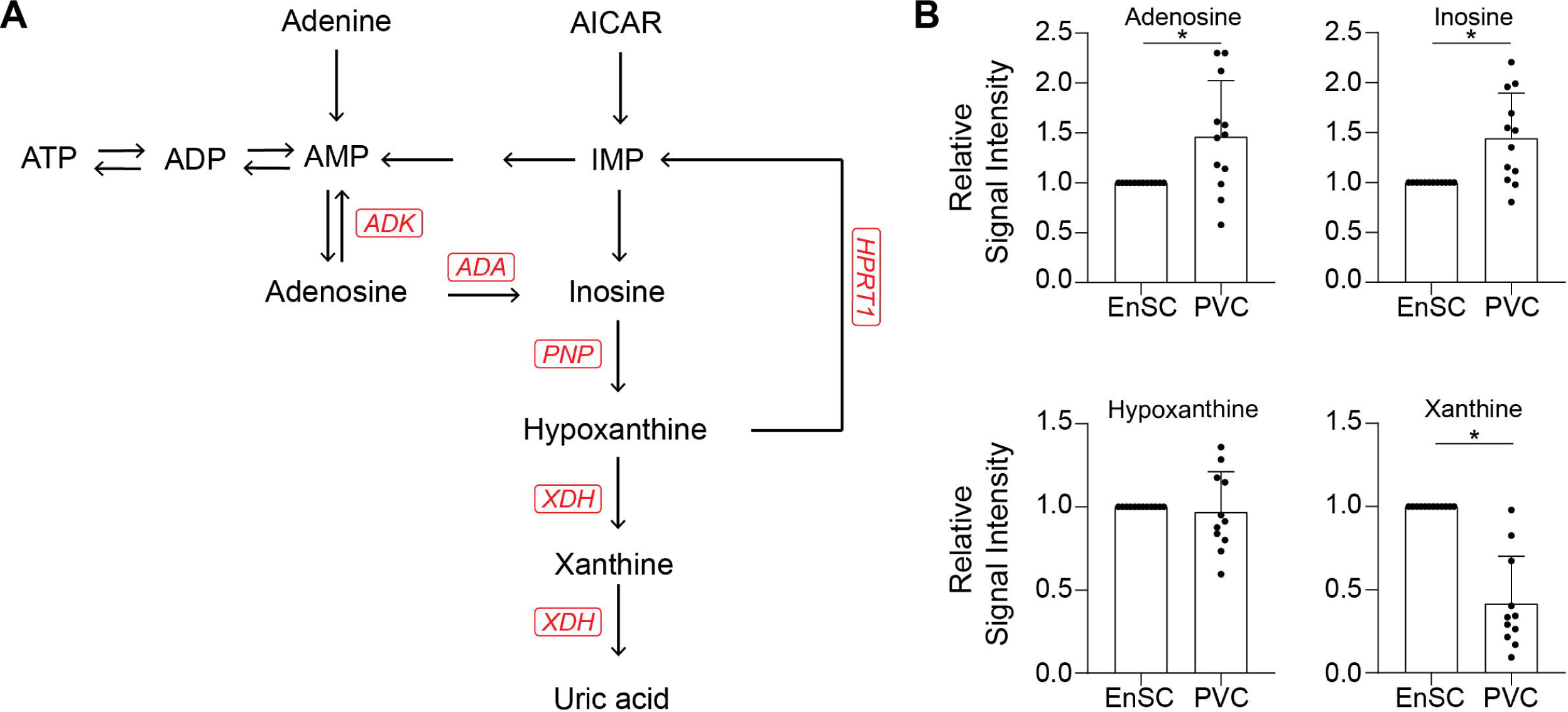
Targeted metabolomic analysis. A) Schematic overview of the purine metabolic pathway. B) Secreted purine metabolites were measured by targeted mass spectrometry in conditioned medium of paired PVC and EnSC cultures decidualized for 8 days. The data show the relative change in secreted metabolite levels in PVC compared to EnSC cultures. The data are mean ± SD and the individual data points show variability between paired cultures. * indicates *p*<0.05 (*t*-test with Holm-Sidak correction).

## DISCUSSION

Several studies have documented the dynamic changes in the EnSC transcriptome, proteome and secretome upon decidualization, ^4,46,47^ but changes in secreted metabolites that may function as signaling molecules have not yet been characterized. Here, we demonstrate that decidualization of EnSC and PVC induces conspicuous metabolic footprints that are tightly regulated in a temporal fashion. For instance, HA secretion was prominent upon full decidualization (day 8) in both EnSC and PVC. This ECM macromolecule has gel properties and is typically highly expressed in rapidly expanding tissues. ^48^ The temporally restricted pattern of HA secretion upon decidualization, therefore, suggests a potential role in decidual expansion in early pregnancy. HA also binds CD44 expressed on extravillous trophoblast and vasculature, indicating a role in spiral artery remodeling. ^45^ Moreover, loss of HA secretion has been linked to miscarriage, ^45^ which further underscores the importance of appropriate temporospatial secretion of this ECM component in pregnancy.

Our untargeted analysis also revealed significant changes in secreted lipid metabolites upon decidualization. Phosphodimethyl-EA and PC (16:0/16:0) are present in phospholipid bilayers, act as signaling molecules, and contribute phosphoethanolamine and phosphocholine, respectively, to form sphingolipids such as sphingomyelin. ^49–51^ Decreasing levels of Phosphodimethyl-EA, PC (16:0/16:0) upon decidualization occurred in concert with increased secretion of glucosylsphingosine, indicating a role for extracellular sphingolipids. Enhanced sphingolipid secretion corroborates recent findings from mice demonstrating that genes encoding sphingolipid producing enzymes are upregulated upon decidualization. ^44^ Mice deficient of these sphingolipid synthesizing enzymes exhibit impaired decidualization, reduced implantation sites, and vascular endothelial defects that compromise trophoblast from invading the maternal vasculature. ^43,44^ Sphingolipids play a role in angiogenesis and promote phospholipase A_2_ enzymatic activity, the enzyme required to form arachidonic acid, a precursor for prostaglandins and leukotrienes. ^52,53^ While 20-trihydroxyleukotriene secretion decreased upon decidualization, 2,3-dinor-8-isoPGF_2α_ levels were consistently elevated across the time-course, suggesting that sustained *de novo* synthesis of sphingolipids is important for decidualization, as demonstrated in mice. ^44^

Exometabolomic differences were also observed between PVC and EnSC. The most notable differences occurred on day 2 of decidualization, i.e. coinciding with the initial inflammatory phase. ^13^ The secreted response at this time-point was also more pronounced in PVC than EnSC, in keeping with the topology of decidualization *in vivo*. ^4^ The PVC exometabolome on day 2 was enriched in higher energy metabolites linked to pyruvate metabolism and TCA cycle, indicative of Warburg effect or active metabolism. ^54^ On day 8 of the decidual time-course, the PVC exometabolome was enriched for more complex metabolites, such as sphingolipids and purines involved in signaling and structural integrity. ^49,55^

Purines, along with pyrimidines, stimulate purinergic receptors, which belong to two subfamilies, the P2X and P2Y receptors (P2XR and P2YR, respectively). P2XR are plasma membrane channels that increase cytosolic Ca^2+^ concentration and mediate the flux of K^+^ and Na^+^ upon activation. ^56^ P2X4R is linked to PGE_2_ signaling. ^57^ P2YR are G-protein coupled receptors that modulate Ca^2+^ mobilization and cAMP signaling in response to either the pyrimidines adenine and uridine, or purines. The affinity for different ligands varies between receptors; for example, P2Y2R preferentially binds UTP, whereas P2Y11R binds ATP. ^58^ Purines modulate cell growth, act as coenzymes, and are a source of energy whereas pyrimidines contribute to phospholipid biosynthesis, glycosylation and detoxification. ^59,60^ Not only do endometrial cells, and particularly epithelial cells, express purinergic receptors but placental trophoblast expresses nearly the entire repertoire of purinergic receptors. ^61–63^ Thus, purine and pyrimidine nucleotides potentially act as metabolic sensors that converge on the purinergic receptors at the placental-decidual interface.

Intriguingly, purine levels correlate with the induction of stress ligands expression, such as the cell surface glycoprotein MHC class I polypeptide-related sequence A (MICA), a ligand for the natural killer group 2D receptor (NKG2D). ^64^ Recently, uNK cells were shown to target and eliminate senescent decidual cells through activation of NKG2D receptors, ^5,15^ a process that is purportedly essential to prevent chronic senescence of the placental-decidual interface in pregnancy, leading to tissue breakdown and miscarriage ^15^.

Targeted LC-MS demonstrated markedly lower levels of xanthine in PVC compared to EnSC, irrespective of decidualization. By contrast, decidualized PVC secreted higher levels of adenosine and inosine. Adenosine is an anti-inflammatory metabolite and stress signal,^65^ and parent to inosine. Inosine, an immunomodulator, is exported from cells through nucleoside transporters when intracellular concentrations are high. ^65^ Extracellular inosine signals through adenosine receptors and is broken down to form the downstream metabolite hypoxanthine. Xanthine is formed during the breakdown of hypoxanthine by xanthine dehydrogenase (XDH). In keeping with our observations, *XDH* transcript levels are significantly lower in PVC compared to EnSC.^4^ Lower xanthine secretion by PVC suggests that higher energy purines are important in the perivascular niche as hypoxanthine is readily available to form inosine, a paracrine signaling molecule that can be recycled back to produce upstream metabolites, such as ATP. Extracellular release of ATP can induce a pro-inflammatory state, regulate the binding activity of estrogen receptors, increase production of reactive oxygen species, and induce metalloproteinase expression. ^66–68^ Furthermore, ATP has been shown to promote IL-8 secretion in endometrial epithelial cells and stimulate decidualization. ^69^ Thus, redirecting hypoxanthine towards the upstream purines may represent a mechanism to heighten decidualization of PVC cells. Taken together, the data suggest that a balance and switch between ATP, a mainly proinflammatory molecule, and adenosine, an anti-inflammatory factor, is required for an effective decidual response.

In summary, this study is the first to map the exometabolome changes in differentiating human EnSC and to highlight the potential of different metabolites to act as autocrine or paracrine signalling molecules as the decidual process unfolds. Although the overall temporal change in metabolic footprints was remarkably consistent between decidualizing PVC and EnSC, differential secretion of specific metabolites not only reflects metabolic differences between stromal subpopulations but also raises the possibility of spatial organisation of metabolic cues at the decidual-placental interface.

## Abbreviations

ADA: adenosine deaminase;
ADK: adenosine kinase;
AICAR: 5-aminoimidazole-4-carboxamide ribonucleotide;
DSC: decidual stromal cell;
EnSC: endometrial stromal cell;
FC: fold-change;
HPRT: hypoxanthine-guanine phosphoribosyl-transferase;
IDO: indoleamine 2,3-dioxygenase;
IGFBP1: insulin-like growth factor binding portein-1;
IMP: inosine monophosphate;
MACS: magnetic activated cell sorting;
MICA: MHC class I polypeptide-related sequence A;
MPA: medroxyprogesterone acetate;
MRM: multiple reaction monitoring;
NKG2D: killer group 2D receptor (NKG2D);
P2R: purinergic receptors;
PGE2: prostaglandin E2;
PNP: purine nucleoside phosphorylase;
PRL: prolactin;
PVC: perivascular cells;
ROS: reactive oxygen species;
SUSD2: sushi domain containing 2;
uNK: uterine natural killer cells;
XDH /XOR: xanthine dehydrogenase / xanthine oxidoreductase

## ACKNOWLEDGEMENTS

We thank participating patients for donating endometrial samples. This study was supported by the National Research Foundation Singapore under its NMRC Centre Grant Programme (NMRC/CG/M003/2017), the National Research Foundation Singapore Fellowship (NRF-NRFF2017-03) to Q.C. and Wellcome Trust Investigator Award to J.J.B. (212233/Z/18/Z). S.H is funded by the University of Warwick Medical School and A*STAR, Singapore as part of the A*STAR Research Attachment Programme.

## CONFLICT OF INTEREST STATEMENT

The authors have stated that there are no conflicts of interest with this article

## AUTHOR CONTRIBUTIONS

Conceptualization: Y.H.L.; Research: J.Z., M.D.C., E.S.L., C.L., K.M.; Resources: Y.H.L., J.J.B., Q.C.; Data analysis: S.H.: Writing S.H., Y.H.L., J.J.B; Supervision: Y.H.L, J.J.B., Q.C.

## Notes

### Competing Interest Statement

The authors have declared no competing interest.

