## Supplementary figures and images for "Exometabolomic analysis of decidualizing human endometrial stromal and perivascular cells"

### Figure S1

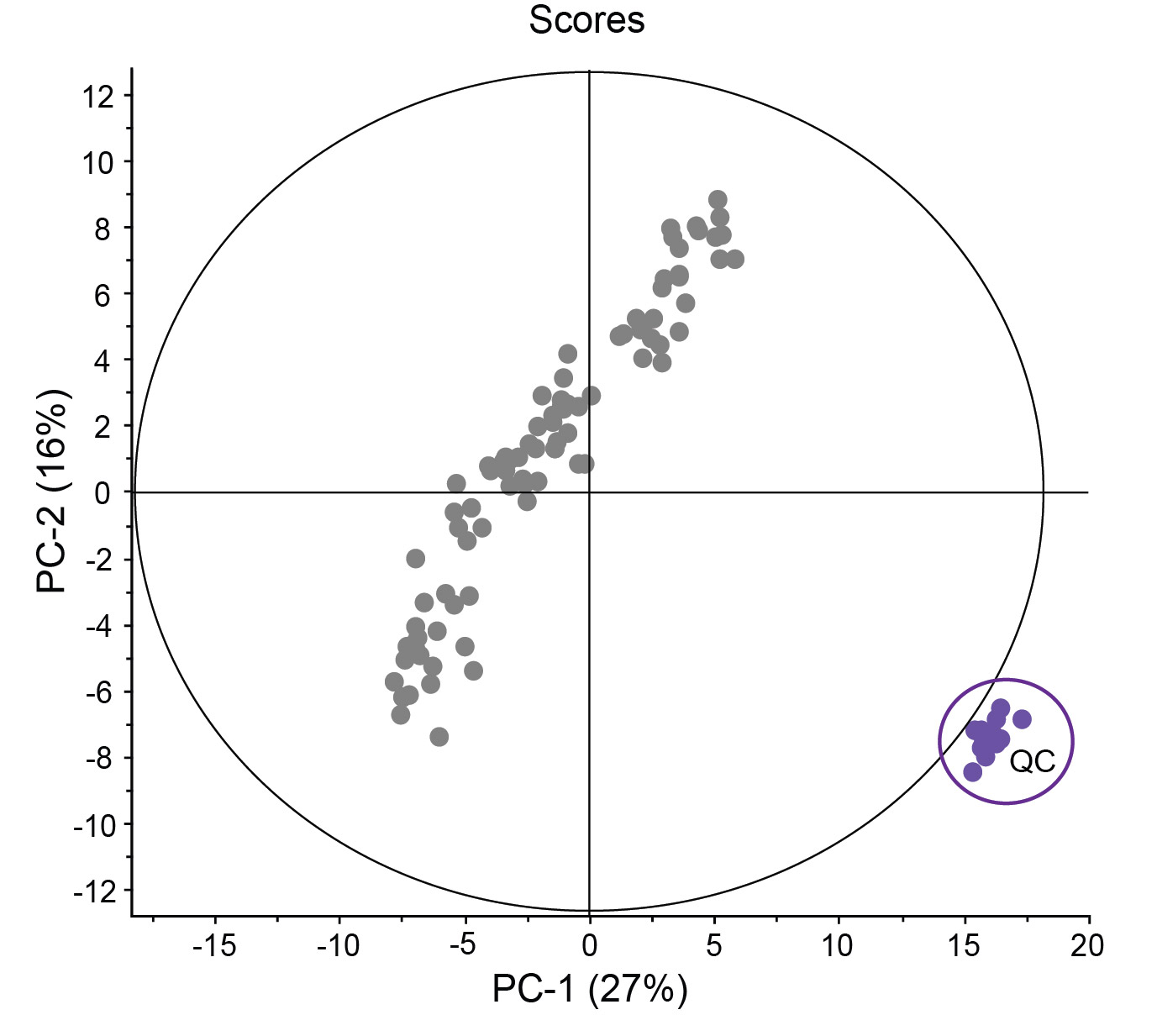

### Figure S2

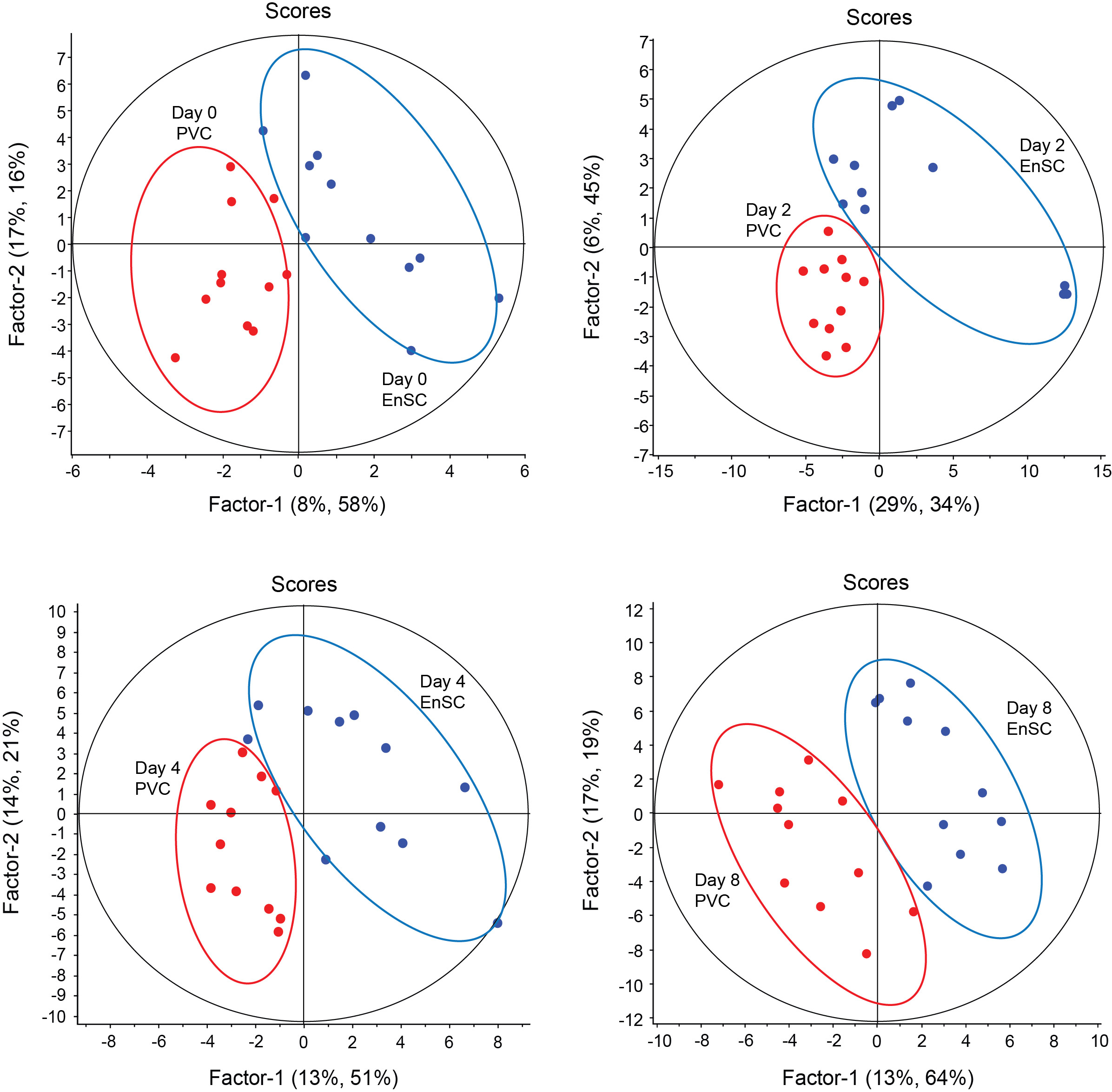
